# Generalized brain-state modeling with KenazLBM

**DOI:** 10.1101/2025.08.10.669538

**Authors:** Graham W. Johnson, Ghassan S. Makhoul, Derek J. Doss, Bruno Hidalgo Monroy Lerma, Leon Y. Cai, Emily Liao, Danika L. Paulo, Anas Reda, C Price Withers, Addison C. Cavender, Helen Qian, Derrick Obiri-Yeboah, Kobina Mensah-Brown, Panagiotis Kerezoudis, Matthew R. Baker, Michael A. Jensen, Shilpa B. Reddy, Shawniqua Williams Roberson, Angela N. Crudele, Robert J. Kahoud, Robert P. Naftel, Maximiliano A. Hawkes, Vaclav Kremen, Mohamad Bydon, Rushna Ali, Kendall Lee, Sarah K. Bick, Nuri F. Ince, Jamie J. Van Gompel, Christos Constantinidis, Victoria L. Morgan, Richard W. Marsh, Gelareh Zadeh, Gregory A. Worrell, Kai J. Miller, Dario J. Englot

## Abstract

The large-scale functional state of a human brain remains difficult to characterize, much less predict. Regardless, techniques have been engineered to electrically neuromodulate the brain to treat a subset of neurologic and psychiatric disorders with moderate efficacy. Accurate characterization of a brain’s instantaneous functional state has stymied the development of more effective neuromodulation paradigms. Advanced computational methods are required to address this gap and enable large-scale neuroscience. Here we define the concept of generalized brain-state modeling across humans as Large Brain-State Modeling (LBM) and present KenazLBM as the world’s first example. KenazLBM can instantaneously characterize the functional state of a person’s brain with raw iEEG data, and predict future brain-states. KenazLBM was trained on over 17.9 billion unique multichannel tokens from people undergoing intracranial electroencephalography (iEEG) recordings, and has learned to interrelate brain-states between people into a common interpretable topology. Most importantly, the model generalizes to unseen subject data with significant recording channel heterogeneity from the training set. We offer KenazLBM as a first generalized brain-state model to serve as a new paradigm of basic neuroscience inquiry and potential translation into neuromodulation therapeutics.

## 1 Main

How do we define the state of a brain? In everyday life, we may observe someone to be sleeping, angry, talking, or focused. Clinically, we may define states such as depressed, ictal, manic, or psychotic. However, these are all extrinsic definitions based on observable behaviors. Neuroscience has long sought an intrinsic quantifiable neurophysiological correlate for these highly abstracted global brain-states. There are multiple modalities to measure brain activity, but the knowledge gap lies in how to digest these data on a massive-scale to characterize brain-state. Furthermore, could a person’s future brain-state be predicted, and potentially modulated? The goal of this work was to develop a paradigm to elucidate highly-abstracted brain-states in a framework that generalizes to unseen subjects. Specifically, we aimed to utilize a presumed fundamental relationship between brain activity across humans to train what we define as a Large Brain-State Model (LBM) capable of instantaneously capturing a new subject’s clinically relevant brain-state and accurately predicting the future brain-state trajectory (Fig. 1). This paradigm is agnostic to clinical disorder and is broadly applicable to any brain-state of interest. The LBM concept draws inspiration from large language models (LLM) and pathology foundation models (e.g. Clinical Histopathology Imaging Evaluation Foundation - CHIEF) and serves as the first example of a generalizable brain-state model.[1, 2]

**Fig. 1.**
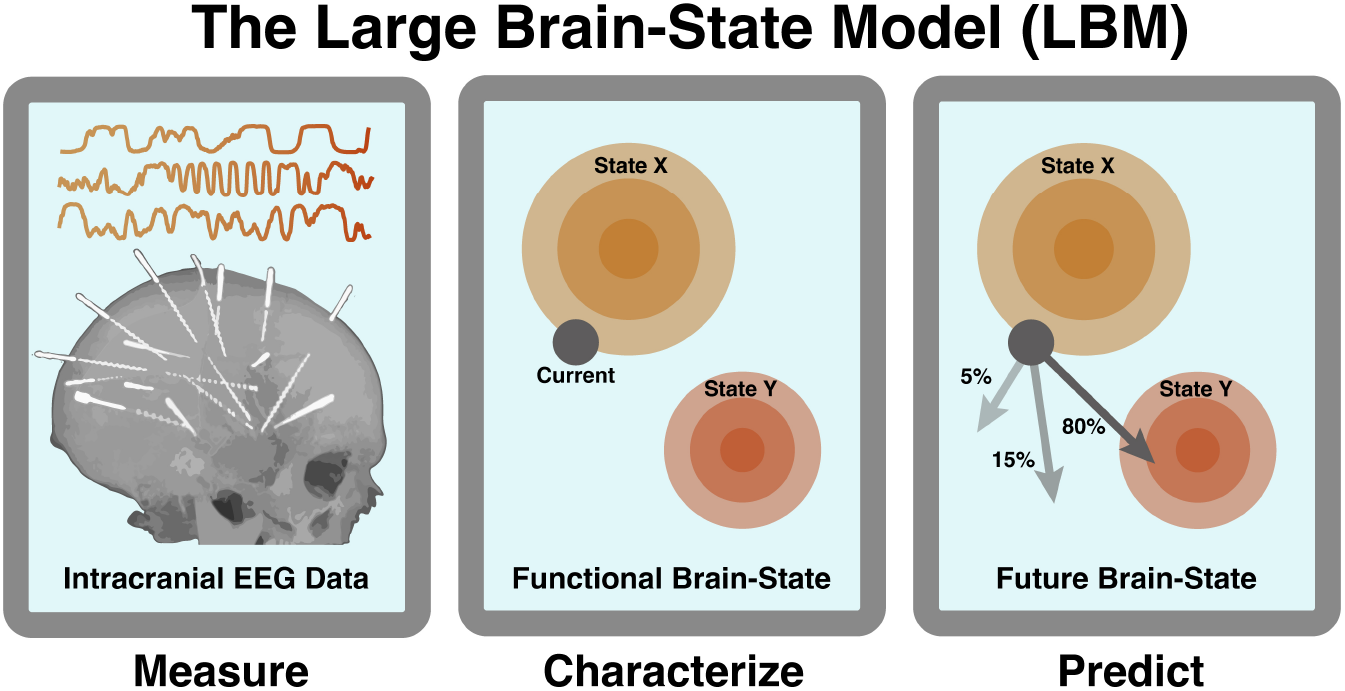
Overview of the Large Brain-State Model (LBM) paradigm. The goal of this generalized framework is to maximally extract biologically-relevant information from iEEG recordings to characterize the current state of a brain and predict future states. This paradigm was designed for direct applicability to adaptive neuromodulation for neurologic and psychiatric diseases. We coin the term LBM to describe the use of a generalized foundation model to drive brain-state characterization, prediction, and modulation.

To develop an LBM, the choice of observable phenomenon was paramount to maximize potential information acquisition and generalization to unseen subjects. Many neurophysiological activity measurement techniques exist at various scales of resolution. For example, in increasing order of abstraction, we have ex vivo patch clamping,[3] in vivo single unit microelectrode recordings,[4, 5] local field potentials (LFP) from intracranial macroelectrodes,[6, 7] neural circuit dynamics/connectivity between LFPs, [8] and whole brain acquisition techniques like functional magnetic resonance imaging.[9] We chose intracranial electroencephalography (iEEG) depth electrode recordings from humans undergoing clinical workup for drug-resistant epilepsy - These data offered the highest temporal resolution with minimal artifact, albeit sparse spatial sampling, and are recorded for days to weeks continuously per subject.[10]

The next design decision was how to embed LFP information from iEEG data into a meaningful high-dimensional brain-state space. We opted for a transformer-based design with a highly randomized training paradigm to augment our 405 days (17.9 billion tokens) of iEEG recordings into virtually infinite input combinations (Fig. 2a-b). The result is a pretrained generalized brain-state model, KenazLBM, that is capable of ingesting unseen subjects’ iEEG data and projecting their brainstate onto an interpretable manifold, and accurately predicting future brain-states. A particularly powerful emergent property of the model is that KenazLBM is agnostic to input channel order, and robust to varying iEEG implant schemes (Fig. 2c). The pretrained weights and a step-by-step guide on how to utilize KenazLBM is available in the Supplementary Info: Appendix A. We anticipate that by providing this framework for open academic use, we can foster collaboration and advancement of this novel paradigm.

**Fig. 2.**
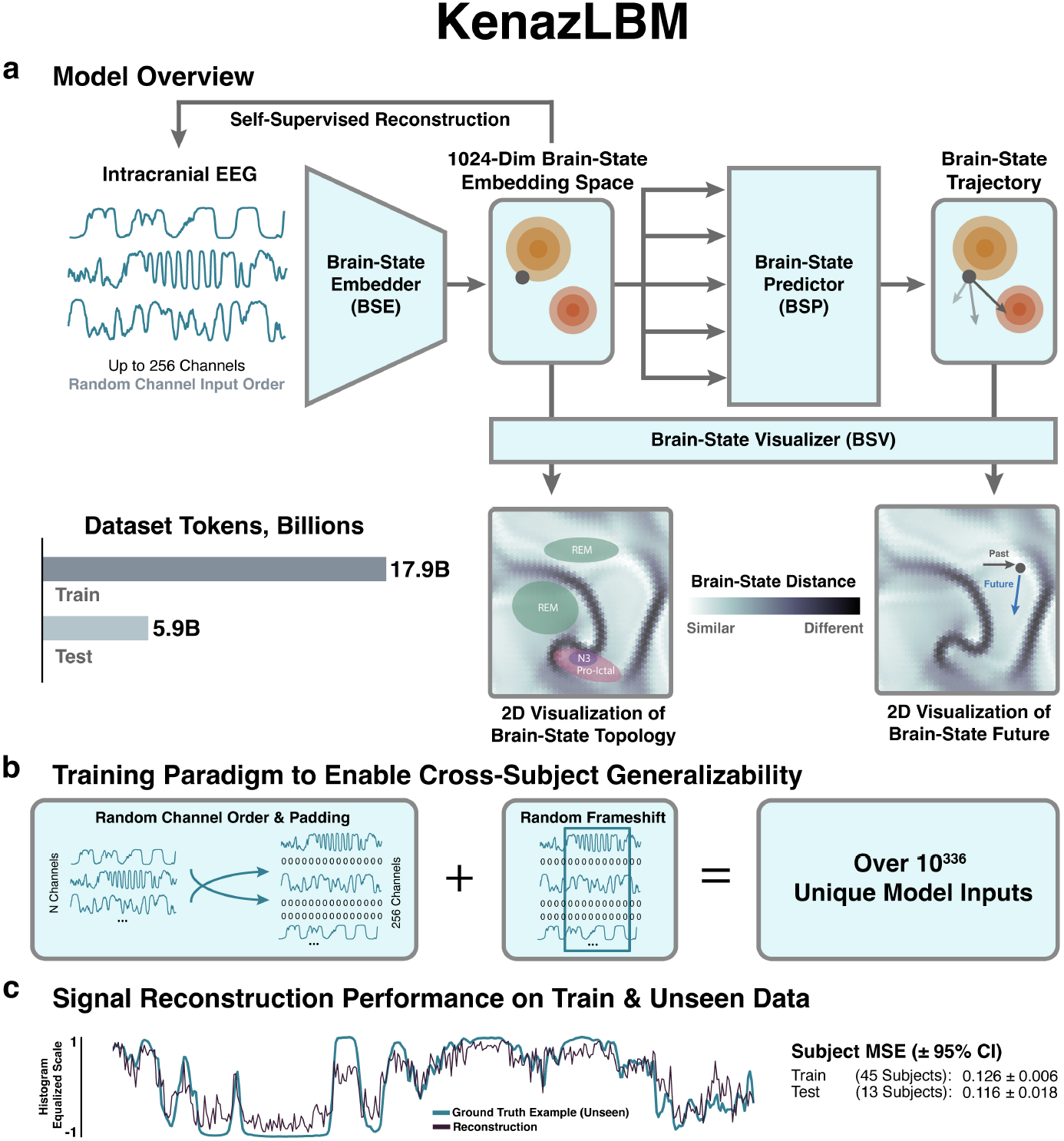
Overview of how data flows through the model and results of generalized raw signal reconstruction. a) iEEG data is first embedded into a common latent space with the Brain-State Embedder (BSE). Next, future embeddings (i.e. tokens) are predicted with the Brain-State Predictor (BSP). The embedding spaces of the BSE and BSP are 1024 dimensions. Thus, to interpret this high-dimensional space, the Brain-State Visualizer (BSV) brings the dimensions down to two and organizes brainstates with a Kohonen Self-Organizing Map (SOM). b) To maximize the model’s ability to generalize to unseen subject data, a heavily randomized training paradigm was used. Specifically, for every forward pass through the model, all subject’s channels were randomly ordered and randomly padded with zeros up to 256 channels. Further, a random frameshift was applied that takes the 17.9B training tokens into virtually infinite input variations into the model. c) The result of this generalized model architecture and training paradigm is indistinguishable reconstruction performance on raw iEEG data reconstruction between training data and unseen test data. MSE = mean squared error.

### 1.1 KenazLBM characterizes interpretable

**clinically-meaningful, and generalized human brain-states**

The KenazLBM architecture operates in 1024-dimensional latent space. Thus, the Brain-State Visualizer (BSV) has been trained in parallel to help interpret the lower-dimensional manifold that exists within the high-dimensional space (Fig. 2). Specifically, a Kohonen Self-Organizing Map (SOM) was built to aggregate similar brain-states.[11] It is important to note that there was no element of supervision in the training process - i.e. all levels of the KenazLBM architecture are unsupervised or self-supervised, including the SOM creation. The SOM of the training and completely withheld test datasets can be seen in Fig. 3. The grayscale colors in Fig. 3a-b represent the “U-Matrix” of the SOM where darker colors indicate brain-states that are further apart in the original 1024-dimensional space. Thus the dark ridges in Fig. 3a-b are akin to peaks in brain-state topology that are unlikely for a brain to traverse. The SOM is of hexagonal geometry and wraps up-down and left-right into a toroidal surface which is flattened into 2D here. Clinically-relevant labels can then be overlaid post-hoc onto the SOM, as is done in Fig. 3a with the 4-hour pre-ictal density.

**Fig. 3.**
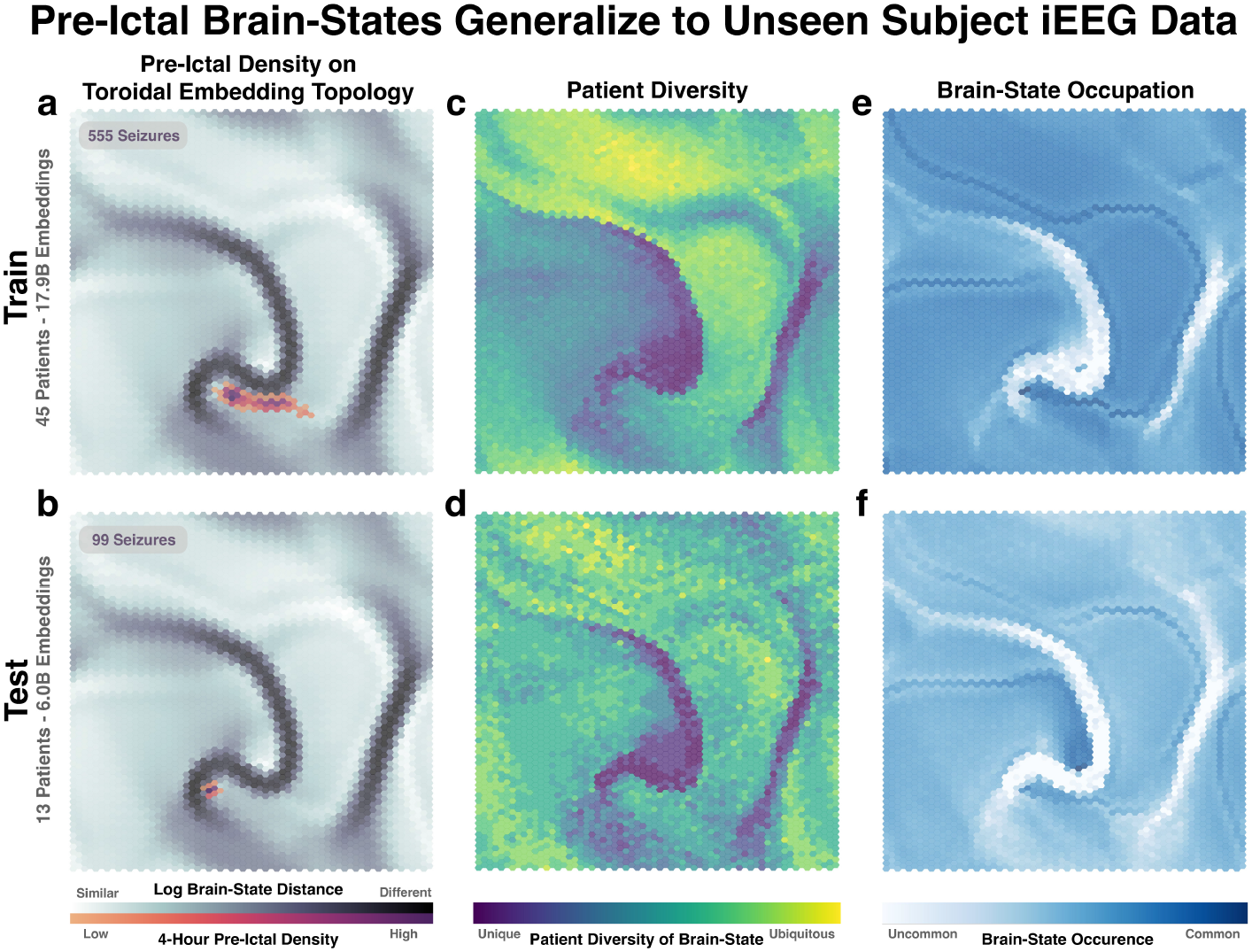
Post-hoc interpretation of KenazLBM embedding space. The KenazLBM model was trained on 17.9B unique input tokens, which comprises over 405 days worth of iEEG data from 45 subjects. The model was then tested on 6.0B tokens which comprises 135 days worth of iEEG data from 13 subjects. Importantly, the test data were completely withheld from model training. The visualizations shown here are dimensionality reductions of the 1024 latent space into 2D with the help of the BSV. The BSV consists of a variational autoencoder and a Kohonen Self-Organizing Map (SOM). The SOM is based on a hexagonal geometry and is wrapped vertically and horizontally into a toroid configuration (i.e. going off the right of the map wraps around to the left and similarly for up and down). The top row of plots represent the training data, and the bottom row represents completely withheld test data. a) The white and gray visualization of the SOM is known as the U-Matrix and represents the distance each brain state is from its neighbors. A darker color on the U-Matrix implies that the brain-state is very far from it’s neighbors on the native 1024 dimension manifold. Thus, black ridges can be thought of as mountain ranges that are difficult for the brain-state to navigate across. The U-Matrix is a static representation of the topology of the brain-state embedding space, and we can then add post-hoc labels to the U-Matrix to represent clinically important states. Here we have added a contour of 4 hours from known pre-ictal events thresholded at a density of 0.5. There were 555 seizures in the training data that were annotated by board-certified epileptologists. We defined preictal as 4 hours before each of these electroclinical seizures. We define areas included in this pre-ictal density and areas adjacent to these areas as “pro-ictal” to indicate high seizure risk states. b) The U-Matrix visualization is the same as above, but now we have overlaid the completely withheld test data 4-hour pre-ictal density onto the plot. As expected with a properly generalized model, the test pre-ictal density aligns with the region we have considered pro-ictal based on the training data. c-d) Yellow indicates states visited by every subject, where purple represents states only visited by a few subjects. e-f) Blue indicates more overall occupancy of a brain-state throughout the entire dataset.

Specifically, we took the 555 epileptologist-defined electroclinical seizures represented across the train dataset subjects and labeled the 4-hour pre-ictal period thresholded at 0.5 contour density. This region of the SOM is what we call “pro-ictal” because there are many instances of subjects visiting this area, but not immediately evolving into a seizure. This is the desired utility of KenazLBM - to aggregate functionally similar brain-states and sparsely label them with incomplete clinical information. Importantly, the pro-ictal region of the SOM generalizes to the completely withheld 13 subjects comprising 99 electroclinical seizures from the test dataset (Fig. 3b). To assess the ubiquity of the brain-states, Fig. 3c-d demonstrate the proportion of patients that visit each brain state - yellow indicates states visited by every subject, where purple represents states only visited by a few subjects. As expected, the states that are black on the U-Matrix (Fig. 3a-b) are not visited by many subjects because they are rare states in general. Finally, Fig. 3e-f represent how often a state is visited. This is different from plots Fig. 3c-d in that a state may be considered diverse if every subject visits that state, but it still may be a very rare state. Thus, Fig. 3e-f showcase how common each state is with darker blue indicating more overall occupancy of that state throughout the entire dataset. As expected, the black ridges in the U-Matrix are rarely visited.

Although these data were gathered from people with drug-resistant epilepsy, the utility of KenazLBM is not restricted to peri-ictal brain-state delineation. To demonstrate the richness of brain-state information present in large-scale iEEG recordings we have overlaid sleep stages as defined by the Montreal Neurological Institute (MNI) SleepSEEG algorithm to demonstrate the brain-state organization of sleep (Fig 4).[12] Of clinical interest, N3 sleep organizes closely to pro-ictal epochs which is in alignment with previous work postulating the relatively high seizure generation risk of N3 sleep. Conversely, rapid eye movement (REM) sleep is across the U-Matrix “ridge” from the pro-ictal brain-state region and is thought to be protective against seizure generation. [13–18] Interestingly, REM sleep is further divided into 2 distinct subgroups which suggests that we are oversimplifying this sleep stage by defining both as REM – which is perhaps explained by phasic vs. tonic REM.[19] Overall, KenazLBM shows promise in defining a new generalizable state space upon which to test hypotheses and develop neuromodulation interfaces.

**Fig. 4.**
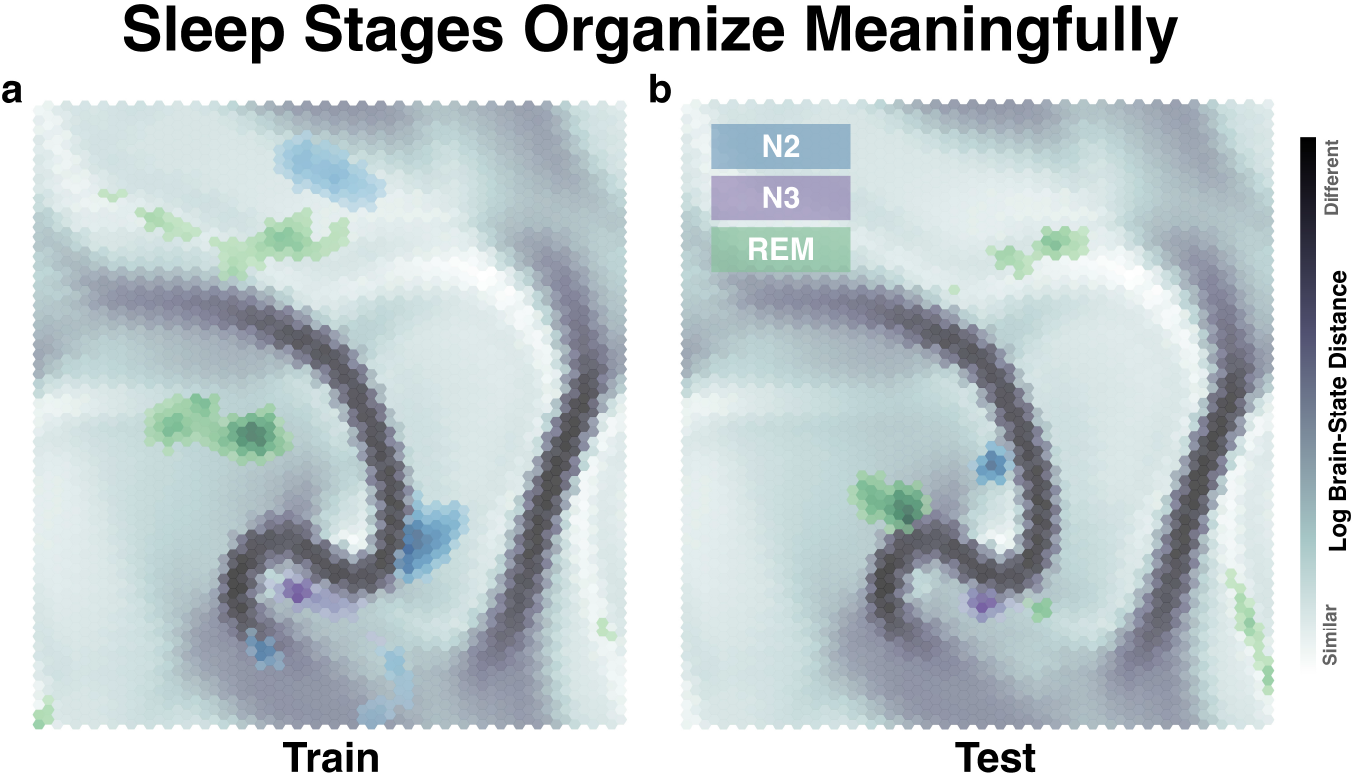
Sleep stages generalize to unseen subject recordings. U-Matrix visualization with sleep stages overlaid (N2, N3, REM). Sleep stages were defined with Montreal Neurologic Institute (MNI) sleep staging algorithm SleepSEEG, thresholded at above 50% model confidence for 5 minutes, excluding N1 and excluding any night upon which a seizure occurred. a) Sleep stages for the training data, notably N3 sleep overlaps heavily with the pro-ictal brain-state locations, and REM sleep is very far across high U-Matrix ridges. This observation is in alignment with previous notions that N3 is a high risk state for seizures and REM is generally protective against seizure genesis. b) test data generalizes well for N3 and REM sleep, but not for the more difficult to characterize N2 sleep.

### 1.2 Future brain-states can be accurately predicted on unseen human data

Of particular interest to our group is the advancement of adaptive neuromodulation paradigms to treat neurologic and psychiatric diseases. The utility of KenazLBM in this endeavor is predicated on the ability to predict future brain-states. As shown in Fig. 5, KenazLBM is able to accurately predict brain-state trajectories with comparable efficacy in the training and completely withheld test sets the mean-squared error (MSE) of 100 predictions is displayed overlying an representative prediction for the training (Fig. 5a) and test (Fig. 5b) datasets. The ability to predict the future from a new subject’s raw data that was not utilized in model training is a non-trivial contribution to the future of adaptive neuromodulation paradigms. We hope that this generalized framework can be leveraged in future technologies.

**Fig. 5.**
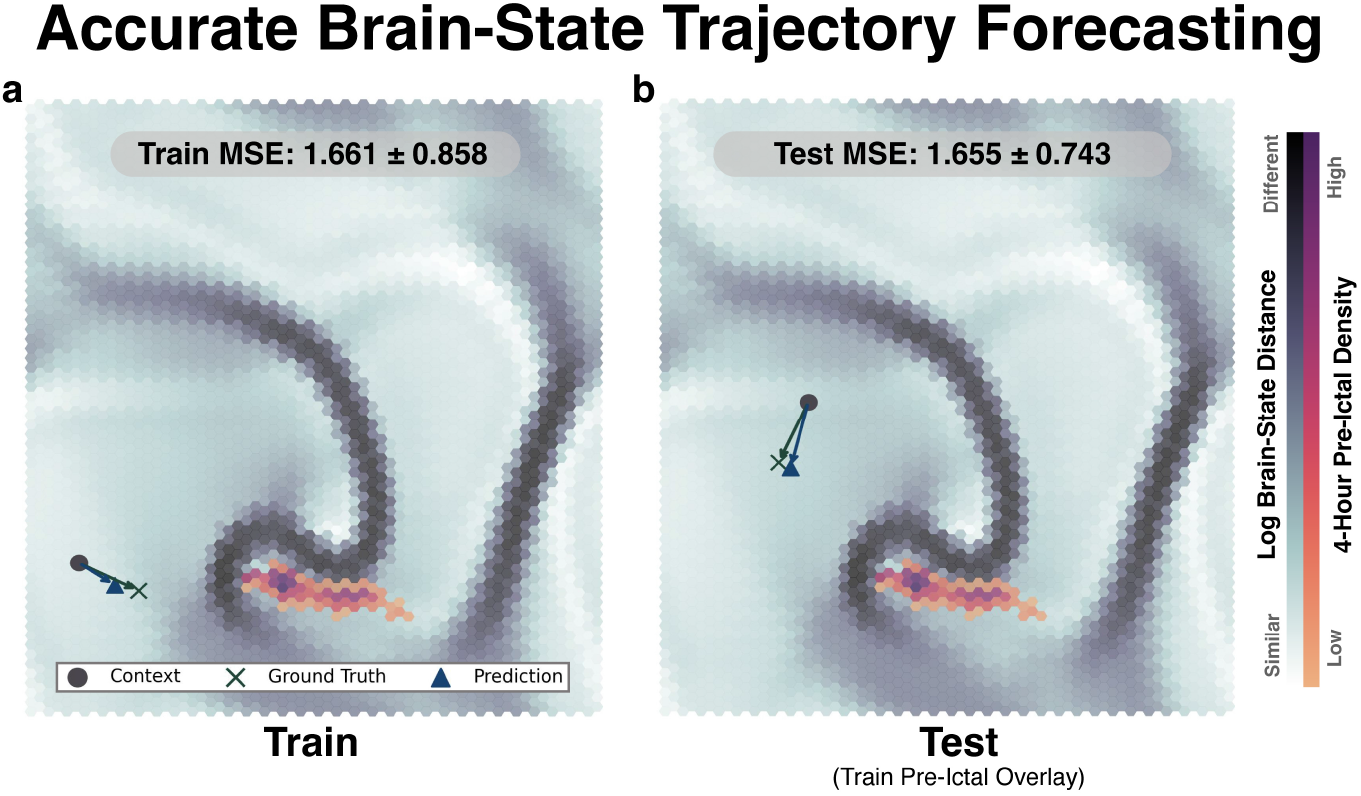
Brain-State trajectory mapping with BSP. A vital concept for possible neuromodulation paradigms using KenazLBM is the ability to predict the future brain-states. Of more significance is the ability to predict future brain-states for subjects not involved in the training of the model. Exemplified here is KenazLBM’s ability to predict future brain-states of subject’s that it is has never seen before. a) An example of trajectory prediction for the training data and for the b) test data. The training mean-squared error (MSE) of the future predictions is indistinguishable from the test data (1.661 ± 0.858 95% CI vs. 1.655 ± 0.743, respectively)

### 1.3 Related Work

A related, but distinct area of work is that of behavior prediction or clinical forecasting models like seizure prediction with EEG.[20–23] KenazLBM was developed in direct response to the pitfalls of such models. Clinical prediction models based on physiologic data are dependent on supervised learning paradigms that leverage expert labeling of clinical data. This technique will always be limited by the quality of the label, the conflation of a single label to potentially distinct states (e.g. REM sleep in Fig. 4), and the confidence of pertinent or assumed negative labels (e.g. all the brain-states adjacent or coincident with the pro-ictal states, but were not labeled as such). Our LBM paradigm does not make any assumptions about data labeling. Thus, KenazLBM serves as a foundation model for brain-state exploration and a source of potential utility in neuromodulation paradigms where a smooth, complete, and continuous manifold upon which to adaptively modulate is crucial.

More similar work to KenazLBM is that of groups utilizing self-supervised frameworks to aggregate brain-states. Specifically, the work of Nejedly et al. that explores the use of a variational autoencoder (VAE) to aggregate similar brain-states in an unsupervised manner on limited epochs.[24] However, as stated above, the majority of similar work in this area is outside of neuroscience due to the limited concentration of machine learning expertise and resources at academic medical centers. Whereas the majority of machine learning expertise and computational resources lie in organizations who develop large-scale models like ChatGPT.[25]

## 2 Discussion

The most relevant areas of discussion are that of KenazLBM’s 1) generalizable properties, 2) interpretation of brain-state space, and 3) potential applicability to neuromodulation.[26–30] To begin, a curious property developed during the highly randomized training of KenazLBM the model seemed to “know” the input channel order to the model. Our first thought was that, despite the virtually infinite input combinations, the model memorized the training subjects’ channels in some manner. However, the proper decoding of channel order was demonstrated even on unseen subjects. That is, the decoder properly organized the output channels to whatever the input order was. Thus, the BSE model seemed to have developed a framework to store a fundamental concept of channel order into the latent space, even for subjects it has never seen before. This is a phenomenon that could yield fundamental insights into the statistical relationship between channels across massive timescale iEEG recordings. Of note, to discourage the Brain-State Predictor (BSP) from harnessing any potential channel information in the latent space, we randomized the channel order for each token in the BSP context forward pass. The result was a BSP that also generalized to unseen subjects by being able to properly predict future brain-states.

Next, the interpretation of brain-state space is only superficially explored in this article. There are a plethora of observable brain-states that could be analyzed including speech, mood, and physical activity. Furthermore, our analyses were limited to that of the 2D SOM representation. Much more extended analyses can be conducted on the higher-dimensional native BSE and BSP latent space. For example, one could explore the properties of continuity, completeness, brain-state loops, and path characteristics. The purpose of our analyses here were to delineate clinically obvious states as an initial validation.

Finally, of particular interest to the health of the human population is that of closed-loop adaptive neuromodulation for the treatment of neurologic and psychiatric diseases. KenazLBM was developed with this particular use case in mind. The simplest question about using an LBM to steer neuromodulation is that of driving “away from bad” states or “toward good” states. To assist this discussion, our group has formulated what we call the “Diseased Brain-State Postulates” (Table 1). These postulates outline how we think about the interpretation of brain-states with the goal of neuromodulating a brain experiencing diseased states. Of particular importance is that of postulate 3 - simply stated, without observing “good” states in a cohort, how can we know “where” in brain-state space to modulate the brain? If the only people receiving iEEG recordings are those with clinically-defined neurologic or psychiatric diseases, how can we obtain high quality physiologic data of healthy brain states? We believe that this is an intractable problem with ethical barriers that will not likely be overcome. Thus, we focus on the concept of neuromodulating “away from bad” states. Much work is to be done to make such a system a reality, but this may be possible with the fundamental insights into the characterization of abstracted brain-states with KenazLBM.

**Table 1.** Diseased Brain-State Postulates with Definitions and Implications used to guide interpretation of Large Brain-State Model embedding spaces.

| Diseased Brain-State Postulates |  |
| --- | --- |
| **Definitions:**<br>• **Embedding space:** The smooth and regularized multidimensional space of possible brain-states constructed from previously observed phenomena (e.g., intracranial electroencephalography). The topology of the high-dimensional embedding space may be visualized with dimensionality reduction techniques onto a two-dimensional map.<br>• **Brain-state/embedding:** The exact mapped location within the embedding space. Brain-state is synonymous with “embedding” in this context.<br>• **Embedding distance:** The mathematical distance between two brain-states in the high-dimensional latent space, as defined by metrics like Euclidean, Manhattan, or angular distance. The higher the embedding distance, the more “different” the brain states.<br>• **Brain-state trajectory:** The sequential path of brain-states occupied through the embedding space for a given epoch of time. |  |
| **Postulate 1** | **No two brain-states antecedent to disease activity are identical.** |
| *[Characterization]* Exact embedding space occupied prior to previous clinically labeled disease events (e.g., seizure, binge-eating, depressive episode) only serves as approximate landmark for future events. Clinically-derived labels may conflate distinct brain states. Therefore, embedding space construction benefits from a purely unsupervised/self-supervised model to allow unlabeled brain-states to aggregate in a meaningful manifold. |  |
| **Postulate 2** | **Temporal evolution of brain-states toward diseased activity can vary.** |
| *[Prediction]* Not all brain-states adjacent to a previously verified pre-disease state will evolve identically – there is inherent stochasticity that is difficult to model. Brain-state prediction models must accommodate this probabilistic nature. |  |
| **Postulate 3** | **Brain-states “far” from observed diseased states are ill-defined.** |
| *[Neuromodulation]* In a subject cohort currently experiencing diseased brain-state activity, few to no “stable” brain states far from disease activity are observed and mapped. Therefore, a theoretical optimal brain-state target for closed-loop neuromodulation may not be readily defined from previously observable brain states. Moreover, there is likely a constellation of potentially stable brain states far from clinically significant disease activity. |  |

Overall, we offer KenazLBM as a first generalized brain-state model to serve as a new paradigm of basic neuroscience inquiry and potential translation into closed-loop adaptive neuromodulation.

## 5 Methods

This work is an evolution from our group’s past work that modeled subject-specific brain-states. [31]. KenazLBM was developed to enable cross-subject brain-state generalization. Thus allowing for one person’s brain-states to inform those of another. And most importantly, obfuscate the need to re-train an entire model from scratch for each subject. All model development was conducted in PyTorch.[32] All pretrained models are freely available under an academic use license at ^***^. Please refer to the supplemental info for a step-by-step guide on how to utilize the pretrained models for new iEEG datasets.

### 5.1 Preprocessing

The training dataset consisted of 45 subjects with iEEG implantation, with a range of 88-190 channels per subject (mean 134.2, std: 26.4). The completely withheld test dataset consisted of 13 subjects with a range of 52-177 channels per subject (mean 127.5, std: 37.3). Channels that were out of the brain parenchyma on post-implantation imaging, and clear artifactual channels were excluded. The raw iEEG data were subsampled to 512 Hz and referenced with an adjacent bipolar montage and filtered with infinite impulse response zero-phase digital filters using pass windows of 1-59, 61-119, 121-179 Hz. To maximize generalization, all subject’s data must be properly scaled into a common range. Further, machine learning paradigms benefit from an equally distributed range of values. Thus, we developed a custom histogram equalization scheme that we call Zero-Centered 1-Dimensional Histogram Equalization (ZHE) (Fig. 6a-c). The ZHE is conducted by reading in the first 24 hours of iEEG recordings for each subject, then parsing the signal into positive and negative values. Each positive/negative domain is processed separately. Each domain is histogram equalized using 10,000 bins. Then each equalized bin is saved as a linear function to be applied to the remainder of that subject’s data (Fig. 6c). The utilization of only the first 24 hours of a subject’s data is what results in an evenly distributed but not perfectly uniform distribution as exemplified in Fig. 6b. This paradigm allows for any new data from that subject to be equalized with an existing scheme without need for an entirely new equalization calculation.

**Fig. 6.**
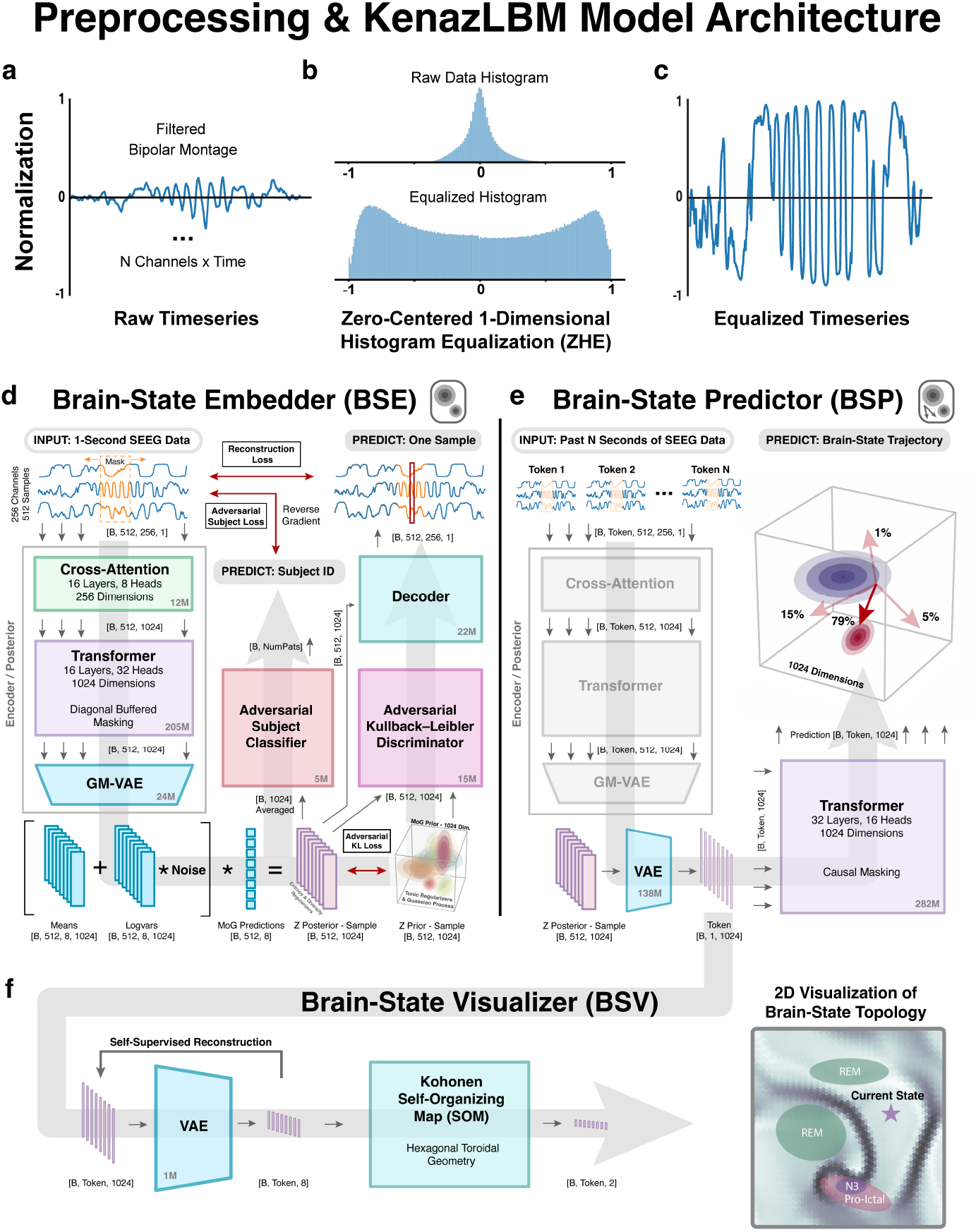
Detailed architecture of the Kenaz Large Brain-State Model (KenazLBM). a-c) The raw iEEG data is first referenced with an adjacent bipolar montage and filtered with infinite impulse response zero-phase digital filters using pass windows of 1-59, 61-119, 121-179 Hz. Next, a custom equalization scheme is used that preserves zero-centering and optimizes the data distribution for machine learning, coined Zero-Centered 1-Dimensional Histogram Equalization (ZHE), and is defined in the Methods section. d) The Brain-State Embedder (BSE) consists of an enocder (a.k.a posterior), decoder, and adversarial training regularizers. For further details refer to the Methods section. e) The Brain-State Predictor (BSP) utilizes the posterior from the BSE. The resampled outputs of the posterior are first fed through a vanilla VAE to reduce the 524,288 redundant BSE embedding space into 1024 dimensions. Next, each token (representing 1 second of iEEG data) is fed into the BSP transformer with context windows of 32 seconds and trained in a traditional manner with causal masking of attention. f) Finally, in order to visualize the BSP embedding space, a Brain-State Visualizer (BSV) is trained that further reduces the 1024 dimensions to 8, then feeds this into a Kohonen Self-Organizing Map (SOM). The SOM is trained on all data smoothed with 64 second windows and a stride of 16 seconds. The SOM aims to properly represent the high-dimensional manifold topology in 2-dimensions.

### 5.2 KenazLBM Architecture

The detailed architecture of KenazLBM can be seen in Fig. 6d-f with data tensor sizes as information flows through the forward pass of the model. The model is split into three distinct modules, The Brain-State Embedder (BSE), Predictor (BSP), and Visualizer (BSV).

#### 5.2.1 Brain-State Embedder (BSE)

There are key components of the BSE architecture and training paradigm that enable the powerful generalization of KenazLBM to unseen subject data. Specifically, the use of a Gaussian-Mixture Variational Autoencoder (GM-VAE) in the BSE to simultaneously promote unique brain-states while also melding subjects’ brain-states into a smooth and complete high-dimensional data manifold in 1024 dimensional embedding space. Next, the randomized training paradigm outlined in Fig. 2b that augments the data input space to virtually infinite permutations and ensures the model does not overfit to certain channel order or temporal event locking. Training stopping criteria was evidence lower bound (ELBO). A powerful result of this paradigm is that the GM-VAE is capable of reconstructing subject data regardless of input order, implantation scheme, or even if the model has seen that subject’s data before. This suggests that the BSE has developed a generalized framework for the fundamental biophysical reality and statistical modeling of iEEG data as a whole.

The BSE (Fig. 6d) consists of an encoder (a.k.a posterior), decoder, and adversarial training regularizers. The first layer of the posterior is a cross-attention block that extracts channel-to-channel motifs. Next, a transformer block is used to extract largescale patterns across time. Importantly, the transformer is diagonally masked with a 16-sample buffer on either side to allow imputation of reconstructed data blinded to 32 samples total. The final layer of the posterior is an 8-component GM-VAE to promote distinct brain-states, but also merge subjects’ brain-states into a smooth and complete manifold in high-dimensional space. As with all VAE paradigms, the output of the GM-VAE is re-parameterized and passed through the decoder to reconstruct a single data point at a time. Next in the forward pass, there are two adversarial components: 1) an adversarial subject classifier with a reversal of gradients during backpropagation to encourage the model to place subjects into the latent space agnostic to subject identification, 2) the Kullback-Leibler calculation for an 8-component GM-VAE is computationally prohibited with Monte-Carlo simulations, thus an adversarial approach is used to train the model to fool a discriminator against samples from the GM 8-component prior. The BSE is trained in isolation to properly balance the delicate adversarial relationship with the intricate GM-VAE regularization.

#### 5.2.2 Brain-State Predictor (BSP)

Next, the BSP is trained in conjunction with the BSV (described below). The pretrained BSE is utilized in inference mode and random channel-order inputs are once again fed into the posterior of the BSE. The outputs of the BSE posterior are then directly fed into the BSP architecture described in Fig. 6e (i.e. the adversarial and decoder components of the BSE are no longer utilized). The first stage of the BSP is a vanilla VAE that takes the redundant 512×1024 dimensional space of the BSE down to 1×1024 dimension tokens where each token represents one second of iEEG data. The BSP then consists of a transformer that is traditionally trained on causal masking of context windows of 32 seconds/tokens of iEEG data to predict next tokens. In parallel, the 1×1024 dimensional space is simultaneously detached from the computational graph and forward passed through the BSV.

#### 5.2.3 Brain-State Visualizer (BSV) with Kohonen Self-Organizing Map (SOM)

The BSV training does not affect the BSP training and is simply a tangential model arm to further reduce the dimensionality of the embedding space for use in clinical interpretation of brain-states. The BSV is a vanilla VAE that takes the dimensions from 1024 to 8. The outputs of the BSV are collected for the entire dataset and used to construct a hexagonal geometry SOM with toroidal wrapping.[11] The SOM is a 64×64 hexagonal grid that wraps left-right and up-down (i.e. a toroid). The entire training and test set are used to construct this visualization. The test set was never used in the BSE, BSP, or BSV training, but it is important to use it in the SOM visualization to allow adjacent locations along the high-dimensional manifold to not be overly simplified by the linear SOM algorithm. The U-Matrix topology best characterizes the high-dimensional manifolds as seen in Fig. 3a-b. Areas of high U-Matrix value correlate to brain-states that are very distant from the nearest neighbor in the hexagonal grid. Thus, the black ridges in the U-Matrix act as mountain barriers that are unlikely to be traversed by a person’s brain-state. Post-hoc clinical labels can then be overlaid to interpret the brain-state topology.

## Supplementary information

Please see online Appendix A for a complete stepby-step guide on how to download and utilize the pretrained KenazLBM on iEEG datasets.

## Declarations

### Ethics approval and consent to participate

All experimental protocols were approved by Vanderbilt Institutional Review Board, and were carried out in accordance with the relevant guidelines and regulations. Informed consent was obtained from all participants and/or their legal guardian.

### Consent for publication

Consent for publication acquired from all parties.

### Data availability

Data available upon reasonable request, see code availability below.

### Materials availability

Not applicable, see code availability below.

### Code availability

All pretrained models and step-by-step instructions on how to utilize are available athttps://kenazlbm.readthedocs.io/en/latest/index.html under academic use license.

### Author contribution

GWJ, GSM, DJD, BHML, LYC, and DJE involved in data curation, project conceptualization, analyses, manuscript drafting, and final revisions. ACD, EL, AR involved in data curation, anlyses, manuscript drafting and final revisions. All remaining authors involved in analyses, manuscript drafting, and final revisions.

## Appendix

### A Step-by-step KenazLBM usage guide

To be disclosed upon publication.

